# State-dependent GABAergic regulation of striatal spiny projection neuron excitability

**DOI:** 10.1101/2023.03.14.532627

**Authors:** Michelle Day, Marziyeh Belal, William C. Surmeier, Alexandra Melendez, David Wokosin, Tatiana Tkatch, Vernon R. J. Clarke, D. James Surmeier

**Author notes:** Correspondence: D. James Surmeier, Department of Neuroscience, Feinberg School of Medicine, Northwestern University, 303 E. Chicago Ave. Chicago, IL 60611 USA,.

## Abstract

Synaptic transmission mediated by GABA_A_ receptors (GABA_A_Rs) in adult, principal striatal spiny projection neurons (SPNs) can suppress ongoing spiking, but its effect on synaptic integration at sub-threshold membrane potentials is less well characterized, particularly those near the resting down-state. To fill this gap, a combination of molecular, optogenetic, optical and electrophysiological approaches were used to study SPNs in mouse *ex vivo* brain slices, and computational tools were used to model somatodendritic synaptic integration. Activation of GABA_A_Rs, either by uncaging of GABA or by optogenetic stimulation of GABAergic synapses, evoked currents with a reversal potential near -60 mV in perforated patch recordings from both juvenile and adult SPNs. Molecular profiling of SPNs suggested that this relatively positive reversal potential was not attributable to NKCC1 expression, but rather to a dynamic equilibrium between KCC2 and Cl-/HCO3-cotransporters. Regardless, from down-state potentials, optogenetic activation of dendritic GABAergic synapses depolarized SPNs. This GABAAR-mediated depolarization summed with trailing ionotropic glutamate receptor (iGluR) stimulation, promoting dendritic spikes and increasing somatic depolarization. Simulations revealed that a diffuse dendritic GABAergic input to SPNs effectively enhanced the response to coincident glutamatergic input. Taken together, our results demonstrate that GABA_A_Rs can work in concert with iGluRs to excite adult SPNs when they are in the resting down-state, suggesting that their inhibitory role is limited to brief periods near spike threshold. This state-dependence calls for a reformulation of the role intrastriatal GABAergic circuits.

## Introduction

The striatum is the largest component of the basal ganglia circuitry regulating goal-directed actions and habits [1,2]. The principal neurons of the striatum are GABAergic spiny projection neurons (SPNs). SPNs integrate information arising from extrastriatal glutamatergic neurons, from intrastriatal GABAergic interneurons and collaterals of neighboring GABAergic SPNs. These intrastriatal GABAergic synapses, which constitute about 20% of all SPN synapses [3], and the post-synaptic GABA_A_Rs transducing the effects of synaptically released GABA, are widely viewed as inhibitory, working in opposition to dendritic excitatory glutamatergic input to suppress SPN spiking [4].

Although the ability of SPN GABA_A_Rs to suppress spiking is clearcut, characterizing them as categorically inhibitory fails to consider two salient features of adult SPNs. First, the commonly described, developmentally regulated, hyperpolarizing shift in the reversal potential of GABA_A_Rs, which transforms GABAergic signaling from excitatory to inhibitory, does not appear to take place in SPNs, at least not to the extent seen in other cell types. Perforated patch recordings from immature SPNs place the GABA_A_R reversal potential near -60 mV – a value close to that seen in other immature neurons [5]. Although this non-invasive recording method has not been applied to adult SPNs, indirect estimates of the GABA_A_R reversal potential have yielded similar values [6,7], suggesting that the developmental shift in the balance of NKCC1 and KCC2 transporters described in other neurons [8] does not happen in SPNs.

Although there does not appear to be a developmental shift in the reversal potential of GABA_A_Rs, other key features of SPN physiology that are relevant to the functional impact of GABA_A_Rs do change in this time window. In particular, SPNs up-regulate the expression of constitutively active, somatodendritic Kir2 K^+^ channels, which leads to a resting membrane potential near the K^+^ equilibrium potential of roughly -80 mV [9]. *In vivo*, this so-called ‘down-state’ in adult SPNs is interrupted by synaptically-driven periods of depolarization when the somatic membrane transitions to potentials near -60 mV, close to spike threshold [10]. Although the synaptic determinants of these ‘up-states’ *in vivo* are not well-defined, *ex vivo* preparations have revealed that the depolarization arising from clustered glutamatergic synaptic activity on distal dendrites can drive regenerative, plateau potentials that mimic up-states [11–13]. A critical trigger of SPN dendritic plateau potentials is the engagement of N-methyl-d-aspartate receptors (NMDARs), which depends not only upon glutamate but membrane depolarization and the displacement of pore-blocking Mg^2+^. This unblocking requires that the membrane depolarize to around -60 mV. Although recent work has shown that dendritic spikes in SPNs can be abbreviated by trailing GABAergic input at the site of glutamatergic stimulation [13], the roles of dendritic location, timing and synaptic strength in determining the interaction between GABA_A_Rs and ionotropic glutamate receptors (iGluRs) have not been systematically explored in SPNs.

To better understand the role of GABA_A_R signaling in adult SPNs, a combination of experimental and computational approaches was employed. These studies revealed that SPNs do not express significant levels of mRNA coding for NKCC1 (at any age) but do express mRNA for KCC2 and HCO3-/Cl-transporters from weaning to adulthood. Surprisingly, given this expression profile, the GABA_A_R reversal potential in SPNs remained near -60 mV from one to nine months of age. Thus, engagement of either synaptic or extra-synaptic GABA_A_Rs excited SPNs in the down-state, pushing them toward spike threshold. Furthermore, both experimental and modeling work demonstrated that dendritic GABA_A_R post-synaptic potentials (PSPs) evoked *prior* to stimulation of ionotropic glutamate receptors effectively worked together to generate a stronger depolarization. Given their sparse spiking *in vivo*, these results suggest that physiological consequences of SPN GABAergic synapses should not be considered as simply inhibitory and that in a wide range of situations GABA_A_Rs work in concert with iGluRs to move SPNs closer to spike threshold and to participate in network function.

## Results

### SPNs robustly expressed mRNA for KCC2, but not NKCC1

GABA_A_R_s_ are primarily permeable to Cl^-^ [14]. Intracellular Cl^-^ concentrations (and hence the equilibrium potential for GABA_A_Rs) are dynamically regulated by a host of transporters, including KCC2, NKCC1 and AE3 [8]. It has been widely hypothesized that the developmental change in the GABA_A_R reversal potential seen in many neurons is governed by a shift in the relative importance of Cl^-^ transporters [8,15]. In immature neurons, the Cl^-^ gradient is thought to be determined largely by NKCC1, which leads to relatively high intracellular Cl^-^ concentration, whereas in adult neurons the gradient is dictated by largely by KCC2, which uses the K^+^ gradient to pump Cl^-^ from the cytosol to the extracellular space leading to lower intracellular Cl^-^ and a hyperpolarized reversal potential [8,15]. To determine whether a similar pattern of expression was evident in SPNs, the striata of 1- and 6-month-old *Adora2*-Cre mice were stereotaxically injected with an adeno-associated virus (AAV) carrying a DIO-RiboTag expression construct [16] **(Fig. 1A)**. Four weeks later, mice were sacrificed; total striatal mRNA and RiboTag-associated mRNA were harvested for quantitative polymerase chain reaction (qPCR) and RNASeq analyses. These experiments revealed that iSPNs robustly expressed mRNA coding for KCC2 (*Slc12a5*), but not NKCC1 (*Slc12a2*) **(Fig. 1B)**; in addition, iSPNs robustly expressed mRNA for the HCO3^-^/Cl^-^ co-transporters AE3 (*Slc4A3*) and NCBE (*Slc4A10*) **(Fig. B)**. The relative expression of these transcripts did not change with development **(Fig. 1B)**. The expression of RiboTag harvested, iSPN-specific transcripts was similar to the mRNA harvested from the entire striatum **(Fig. 1B)**, suggesting that the expression pattern was similar in dSPNs (iSPNs and dSPNs are the principal neurons of the striatum, constituting roughly 90% of all striatal neurons).

**Figure 1.**
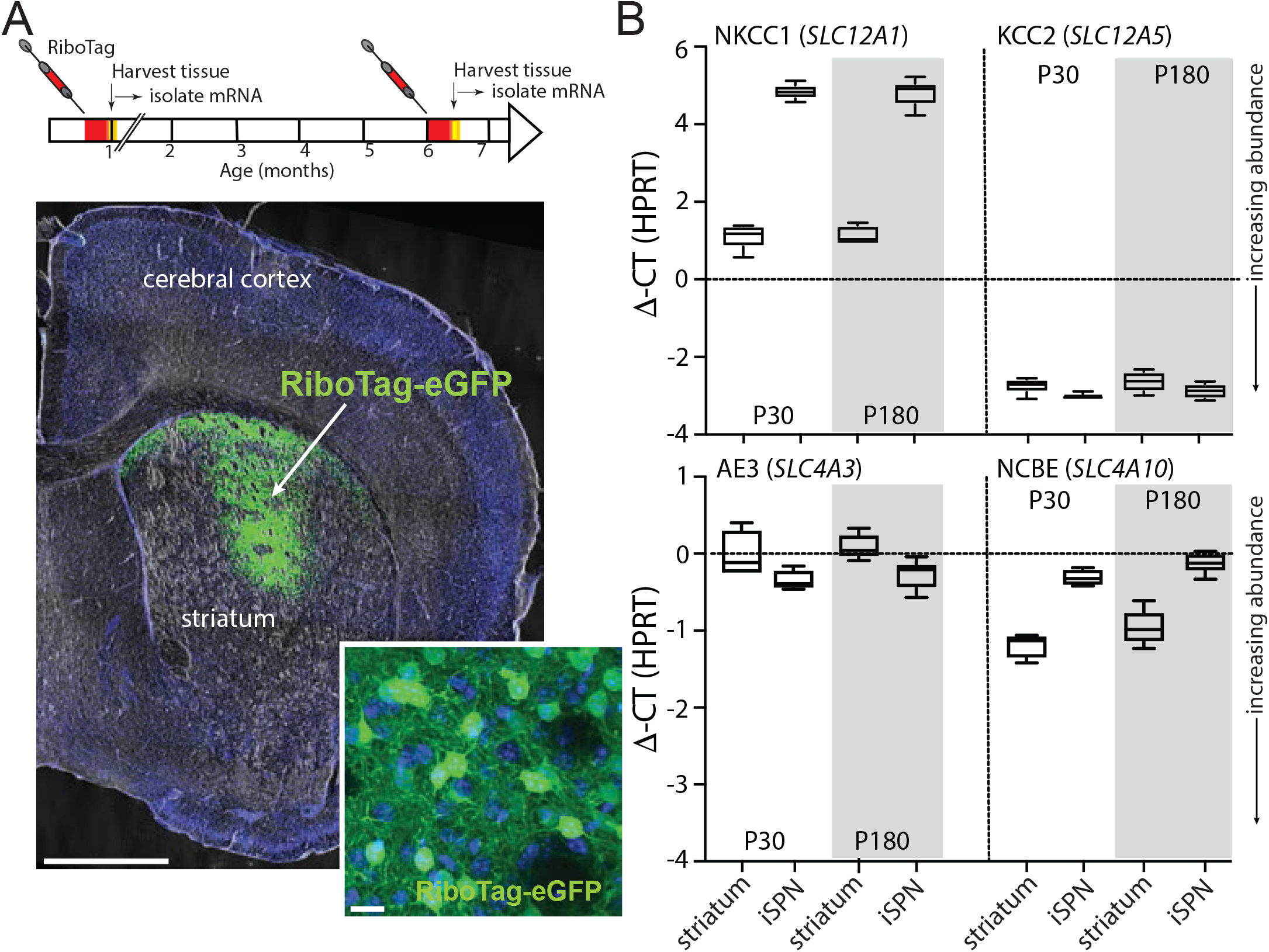
K CC2 mRNA is robustly expressed in SPNs, but not NKCC1. **(A)** The RiboTag construct AAV5-DOI-hSyn-RpI22I1-3Xflag-2A-eGFP was injected into the striatum of Adora2a-cre mice at P18 or at 6 months of age. The coronal slice images demonstrate both the coverage and restriction to the striatum of the stereotaxically-injected AAV carrying the RiboTag and eGFP genes (stereotaxic injection coordinates: ML = -1.85, AP = +0.74, DV = -3.50). Scale bars = 1mm and 20 µm. Ten days later, the infected tissue (green fluorescence) was dissected out with the aid of fluorescence microscopy and qPCR was performed. **(B)** mRNA abundance (DCT) levels for the chloride cotransporters NKCC1 (*SLC12A1*) and KCC2 (*SLC12A5*), along with Cl^-^/HCO_3_ exchangers AE3 (*SLC4A3*) and NCBE (*SLC4A10*), were determined by qPCR in striata from Adora2a-cre mice 4 weeks and 6-months of age.

### The GABA_A_R reversal potential was near -60 mV in both young and adult SPNs

Given the absence of a developmental shift in the expression of genes linked to Cl^-^ homeostasis, our hypothesis was that the GABA_A_R reversal potential in adult SPNs would be similar to the reversal potential reported for immature SPNs [5,6]. To test this hypothesis, *ex vivo* brain slices were prepared from young adult (6-7 months-old) mice and then gramicidin perforated patch recordings were made from identified SPNs. Gramicidin is selectively permeable to monovalent cations, leaving the intracellular Cl^-^ concentration ([Cl^-^]_i_) unperturbed. To visualize dendrites, SPNs were sparsely labeled using an adeno-associated virus (AAV) carrying a SuperClomeleon expression plasmid [17] **(Fig. 2A)**. To activate GABA_A_Rs, Rubi-GABA was uncaged on the soma and dendrites using a blue laser spot **(Fig. 2A)**. The somatic membrane potential was clamped at membrane potentials between -50 and -80 mV prior to uncaging GABA and the resulting currents monitored. The amplitude and polarity of uncaging evoked currents were then plotted as a function of somatic membrane potential. The estimated reversal potential for somatic GABA_A_Rs was near -60 mV **(Fig. 2C)**. Dendritic uncaging of GABA evoked currents which reversed in polarity with somatic membrane potentials slightly more depolarized than -60 mV **(Fig. 2C)**. The modest deviation in these measurements from those where somatic GABA_A_Rs were engaged probably reflects an imperfect space clamp, rather than any significant intracellular gradients in Cl^-^ concentration [18].

**Figure 2.**
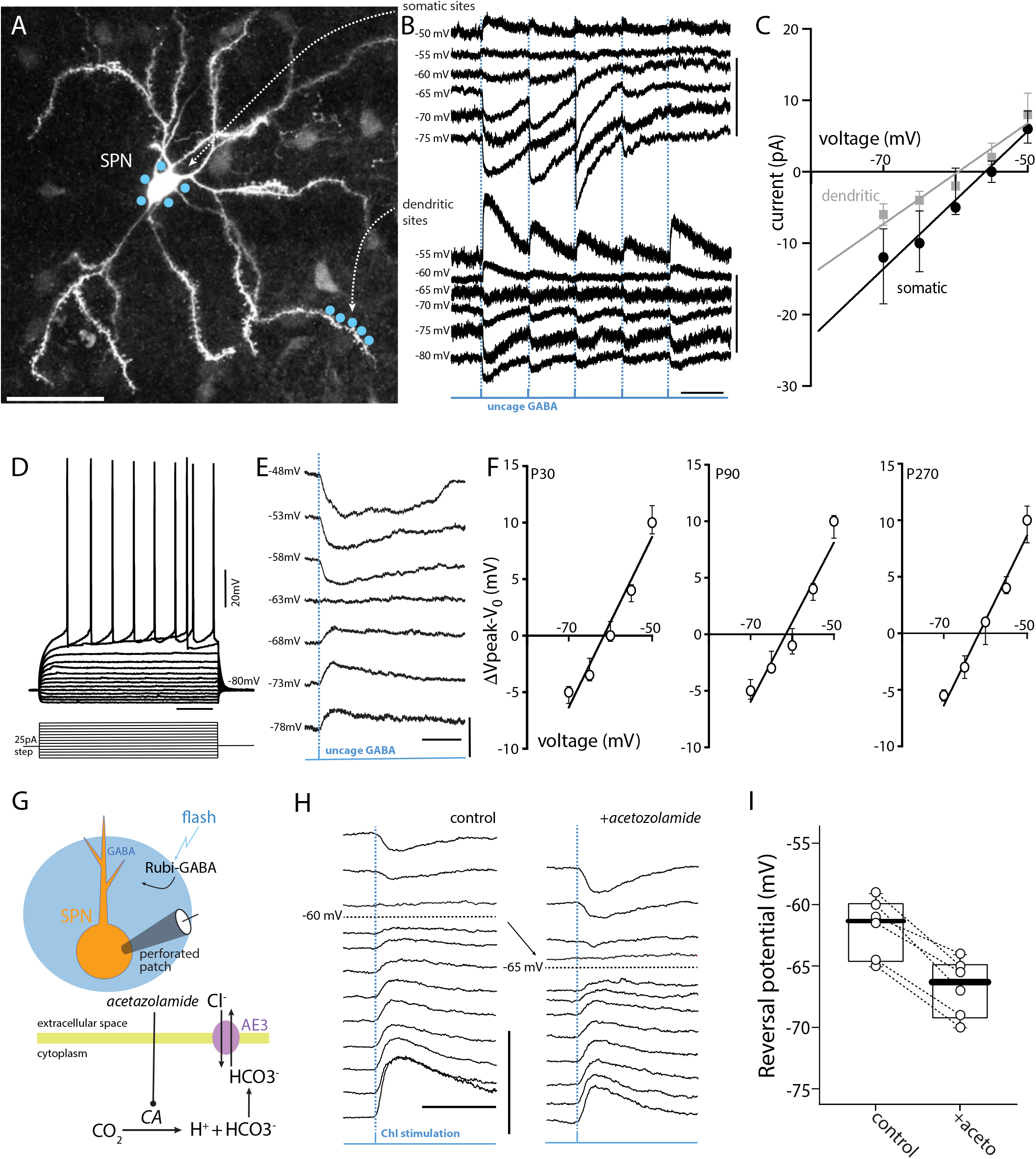
The GABA_A_R reversal potential is near -60 mV in both young and adult SPNs. **(A)** Clomeleon-expressing iSPNs allowed visual identification of dendrites in gramicidin perforated-patch recording conditions where cells cannot be loaded with dyes via internal-pipette solution (920 nm laser maximum projection image, scale bar = 40 µm). When low (<100 MΩ) access-resistance was achieved in voltage-clamp mode, RUBI-GABA (10 µM) was uncaged with a 473 nm laser spot (∼1 µm diameter, 1 ms) in the presence of the synaptic blockers: TTX (1 µM), AP5 (50 µM), NBQX (5 µM), CGP-55845 (1 µM). The laser was targeted to the somatic region or to distal dendrites (blue spots, projection image). **(B)** Representative voltage traces showing GABA responses, recorded in serial, from the soma (top traces, scale bars= 20 pA/2 sec) or the dendrite (lower traces, scale bars = 10 pA/2 sec) as the membrane was manually stepped from -80 mV to -50 mV. **(C)** Plot of the current/voltage relationship between the somatic (black line) and the dendritic (grey line). The data, represented by medians with interquartile ranges, did not differ significantly between the soma and dendritic compartments (n=7). **(D)** Current-clamp experiments in gramicidin perforated-patch mode were performed to examine age-dependent shifts in reversal potential. Here, Adora2a-eGFP positive iSPNs could be visually identified and patched. When low (<100 MΩ) access-resistance was achieved in current-clamp mode, the resting membrane potential along with series of hyperpolarizing and depolarizing steps were used to examine cell health (traces, scale bars = 20 mV/200 ms). **(E)** RUBI-GABA (10 µM) was uncaged over the full-field (3 ms duration, 60x lens) with a 473 nm LED in the presence of the synaptic blockers: AP5 (50 µM), DNQX (5 µM), CGP-55845 (1 µM). Representative current traces showing GABA responses as the membrane was manually stepped from -80 mV to -50 mV, scale bars = 5 mV/200 ms. **(F)** Plot of the change in voltage (ΔV-Vo) at P30, P90, and P270. The data, represented by medians with interquartile ranges, did not differ significantly between the 3 ages tested (n=5 each group). The data shows that the reversal for GABA-induced chloride current is a full 20 mV+ above the resting membrane potential for SPNs, typically, -80 to -85 mV. **(G)** Cotransporters make a significant contribution to the relatively depolarized GABA_A_R reversal potential in SPNs. To assess their contribution to the Cl^-^ gradient, perforated patch recordings were obtained from Adora2a-eGFP iSPNs and then the reversal potential of GABA_A_Rs determined before and after inhibition of carbonic anhydrase (CA) with acetazolamide (AZ, 20 µM). CA catalyzes the conversion of mitochondrially generated CO_2_ to H^+^ and HCO_3-_, which serves as a substrate for HCO_3-_/Cl^-^ cotransporters. **(H)** Representative traces recorded from a visually identified iSPN from an Adora2a-eGFP mouse in gramicidin perforated patch in current-clamp mode in the synaptic blockers: NBQX (5 µM), AP5 (50 µM), CGP-55845 (1 µM), MPEP (1 µM), and CPCCOEt (50 µM). Rubi-GABA (15 µM) was uncaged using a single LED pulse (470 nm, 25 ms). The pulse was applied at an interval of 30 seconds while manually stepping the cell to different potentials from -80 to -50 mV, scale bars = 10 mV/100 ms. **(I)** Summary data shows that application of AZ shifts the reversal of the GABA-induced Cl^-^ current to more negative potentials (p=0.0313, two tailed Wilcoxon sign rank, paired, n=6).

To verify an absence of a developmental shift in the SPN GABA_A_R reversal potential, gramicidin perforated patch recordings were made from SPNs in *ex vivo* brain slices taken from mice at three ages: young (∼1 month-old), young adult (6-7 months-old) and adult (∼9 months-old) mice. SPNs recorded in this mode displayed the characteristic inward rectification, delayed time to the first spike at rheobase, and sustained repetitive spiking with suprathreshold current injection (**Fig. 2D**). As predicted, the membrane potential changes evoked in SPNs by Rubi-GABA uncaging on the peri-somatic membrane reversed near -60 mV at all ages (**Fig. 2E, D**).

Given the low or absent expression of NKCC1 in SPNs, why is the reversal potential of the GABA_A_Rs relatively depolarized? The striatal circuitry is largely quiescent in the *ex vivo* brain slice, making it highly unlikely that ongoing GABAergic signaling was loading neurons with Cl^-^ and pushing the reversal potential in a depolarized direction. Another determinant of the GABA_A_R reversal potential is intracellular HCO_3-_ concentration ([HCO_3-_]_i_) [8]. The intracellular balance between [HCO3^-^]_i_ and [Cl^-^]_i_ is controlled by the activity of plasma membrane HCO_3-_/Cl^-^ cotransporters, like AE3 and NCBE, which, as noted above, are both robustly expressed by SPNs. To assess the role of intracellular HCO_3-_ to the GABA_A_R reversal potential, perforated patch recordings were obtained from SPNs in *ex vivo* brain slices (as described above) and then the reversal potential of GABA_A_Rs determined before and after inhibition of carbonic anhydrase (CA) with acetazolamide [8]. CA catalyzes the conversion of mitochondrially generated CO_2_ to H^+^ and HCO_3-_, which serves as a substrate for HCO_3-_/Cl^-^ cotransporters. RiboTag/RNASeq analysis revealed that iSPNs expressed three different CA subtypes (*Car11>Car12>Car2*) (data not shown). Non-specific inhibition of these CAs with acetazolamide led to a significant negative shift in the reversal potential of GABA_A_Rs (**Fig. 2G, H**), consistent with the proposition that intracellular HCO3^-^ was contributing to the relatively depolarized GABA_A_R reversal potential in SPNs.

### GABA_A_R engagement depolarized SPNs in the down-state

To study the role of synaptically released GABA, a mixed population of striatal GABAergic interneurons were activated by optogenetic stimulation of cholinergic interneurons (ChIs) [19]. Working through nicotinic acetylcholine receptors (nAChRs), ChIs can activate both neurogliaform interneurons (NGFIs) and tyrosine hydroxylase interneurons (THIs) giving rise to GABA_A_R-mediated currents in SPNs [20]. To monitor evoked responses in SPNs, perforated patch recordings were made from identified iSPNs or dSPNs using the approach described above. To ontogenetically-activate ChIs, an AAV carrying a Cre recombinase dependent expression construct for Chronos was injected into the striatum of transgenic mice expressing Cre recombinase under the control of the choline acetyltransferase (ChAT) promoter (**Fig. 3A**). In the *ex vivo* brain slice, SPNs are quiescent and reside in the down-state near -80 mV [10]. As predicted from the GABA uncaging studies above, optical stimulation of ChIs in the presence of ionotropic glutamate receptor antagonists evoked depolarizing, postsynaptic potentials (PSPs) in SPNs that were blocked by the GABA_A_R antagonist gabazine **(Fig. 3B, C)**.

**Figure 3.**
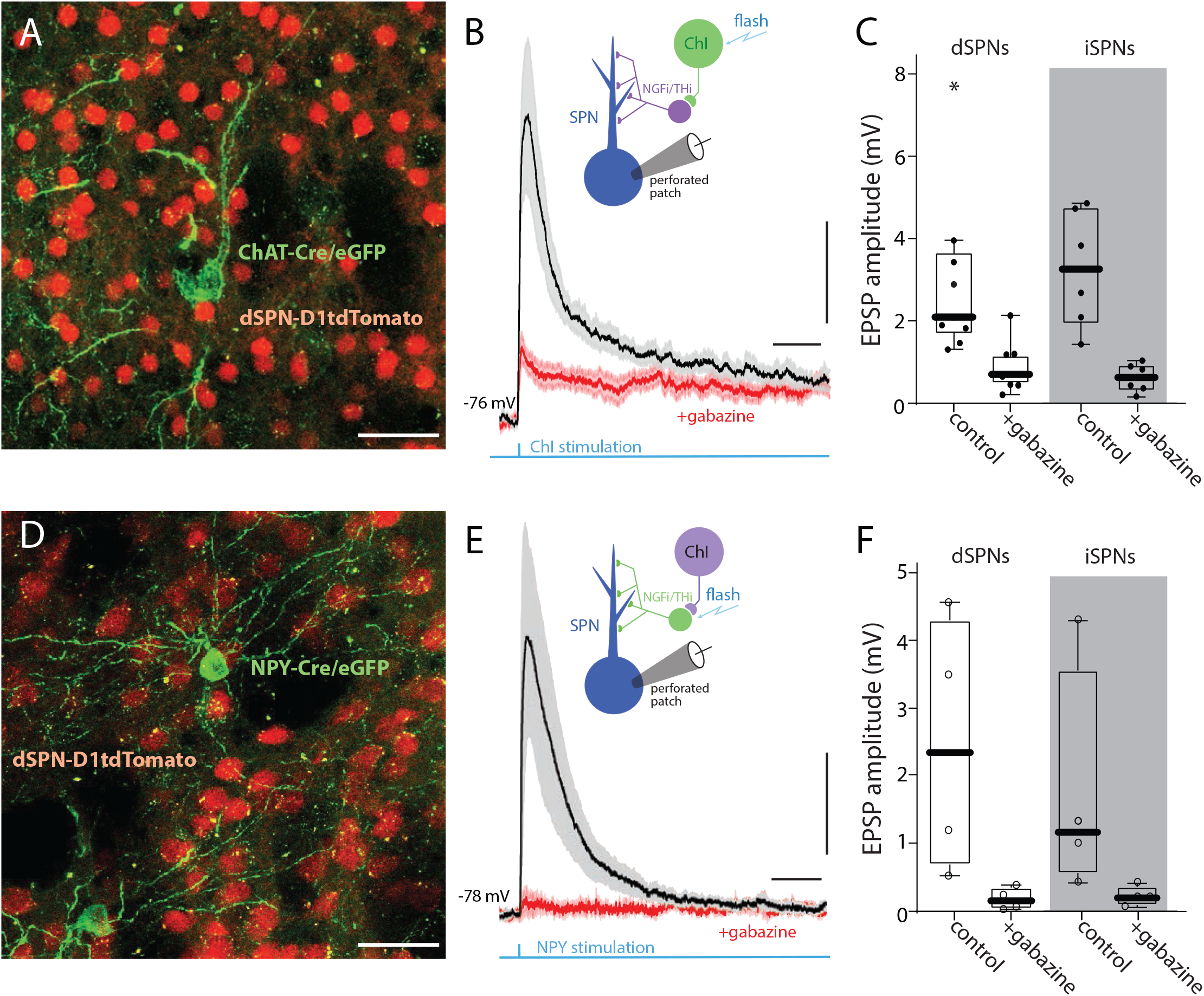
Optogenetic stimulation of ChIs or NPY-expressing interneurons evokes robust EPSPs in both iSPNs and dSPNs. **(A)** AAV9-hSyn-chronos-flex-eGFP was stereotaxically injected to the striatum of two-month-old ChAT-Cre X D1tdTomato mice (Stereotaxic coordinate injection: ML= -1.2, AP=-0.7, DV= -3.4). The coronal confocal slice image shows the expression of Chronos (green cells) in a ChAT-cre neuron (cholinergic interneurons) along with dSPNs expressing tdTomato (red cells, scale bar = 40um). The tissue was dissected and recorded from 21 days post injection. **(B)** The mean average ± standard error of ChI-evoked EPSP responses recorded from visually identified SPNs in gramicidin perforated patch in current-clamp mode in the presence of synaptic blockers: NBQX (5 µM), AP5 (50 µM), CGP-55845 (1 µM), MPEP (1 µM), and CPCCOEt (50 µM). The LED pulse (470 nm, 5 ms) was applied at an interval of 60 seconds. The traces recorded before and after the addition of Gabazine (10 µM). Scale bars = 1 mV/100 ms. **(C)** Summary data shows the median ± interquartile range and the whiskers represent the outer-quartile (dSPNs, n=9) and for (iSPNs, n=6). (**D)** NPY-Cre X D1tdTomato mice were injected as described in *A*. Confocal image showing NPY-Cre neurons expressing Chronos (green) and dSPNs expressing tdTomato (red, scale bars = 40 µm). **(E)** The mean ± s.e.m of NPY-Cre-evoked EPSP responses recorded from visually identified dSPNs in gramicidin perforated patch in current-clamp mode in the presence of blockers as described in *B* before and after the addition of Gabazine (10 µM). Traces from dSPN recorded in NPY (n=4). Scale bars = 1 mV/100 ms. **(F) S**ummary data for dSPNs (n=4) and for iSPNs (n=4).

To simplify the afferent circuitry engaged in these experiments, mice expressing Cre recombinase under the control of the NPY promoter (NPY-Cre) were injected with the same AAV vector used in the ChAT-Cre mice (**Fig. 3D**). NPY is expressed by NGFis and low-threshold spike GABAergic interneurons (LTSIs) [4] – both of which make GABAergic synapses primarily on SPN dendrites [21]. Optogenetic activation of NPY-expressing interneurons alone produced PSPs that were kinetically similar to those evoked by optogenetic stimulation of ChIs **(Fig. 3E, F)**.

### GABA_A_R activation enhanced the depolarization produced by glutamate receptors

As shown previously, in both SPNs and pyramidal neurons [5,6,18], a depolarizing GABA_A_R input can boost the response to a trailing intrasomatic current injection and enhance the probability of spiking. However, GABA_A_R activation during a period of repetitive spiking can suppress spike generation by shunting injected current and by pushing the membrane potential below spike threshold, which is typically between -45 and -50 mV [18].

How might the interaction between GABA_A_Rs and ionotropic glutamate receptors (iGluRs) play out in dendrites? A key feature of SPN dendrites beyond about the first major branch point (∼80 µm from the soma) is the ability to generate dendritic spikes or plateau potentials that can last for 50-200 msec [11–13,22]. These dendritic spikes require the temporal convergence of 10-15 glutamatergic inputs over a relatively short stretch (∼20 µm) of dendrite, which produces enough of a local depolarization to engage N-methyl-D-aspartate receptors (NMDARs) and voltage-dependent Ca^2+^ channels. Previous experimental and modeling work has shown that opening GABA_A_Rs near the site of glutamatergic stimulation after spike initiation can truncate them, much like somatic situation described above. Indeed, as modeling suggests that the dendritic membrane potential during these spikes rises close to 0 mV, GABA_A_R opening should hyperpolarize the dendrites [13].

But, what if the GABA_A_R activation *precedes* the glutamatergic input to dendrites? A priori, one might predict that the dendritic depolarization produced by GABA_A_R opening would enhance the response to trailing glutamatergic input, much like the situation described at the soma. To test this hypothesis, two sets of experiments were performed. Identified iSPNs or dSPNs were recorded from in whole cell mode to allow them to be filled with a dye (Alexa 568) and imaged using two-photon-laser-scanning microscopy (2PLSM) [23]. The [Cl^-^] in the pipette solution was adjusted to yield a GABA_A_R reversal potential near -60 mV. Next, a region of parfocal dendrite was identified to allow two-photon uncaging of DNI-glutamate at visualized spine heads [11,24,25]. In the first set of experiments dendritic GABA_A_Rs were activated by optogenetically stimulating ChIs as described above. Because of their large axonal field and those of the NGFIs/THIs they activate [4,26], optogenetic stimulation of ChIs should produce a diffuse GABAergic input to the dendrites of the recorded SPN **(Fig. 4B)**. As shown above, optogenetic stimulation of ChIs alone evoked a consistent but modest somatic depolarization **(Fig. 4C, D)**. Dendritic uncaging of glutamate alone also evoked a somatic depolarization. The number of axospinous sites stimulated was adjusted to be sub-threshold for dendritic spike generation (assessed by the decay of membrane potential after termination of uncaging) **(Fig. 4C, D)**. When this uncaging event was preceded by ChI-evoked GABA_A_R depolarization, the resulting magnitude and duration of the somatic depolarization was significantly increased in both types of SPN **(Fig. 4C-F)**. Thus, transiently opening dendritic GABA_A_Rs produced a dendritic membrane potential change that enhanced the ability of subsequent dendritic glutamatergic input to push SPNs toward the local spike threshold.

**Figure 4.**
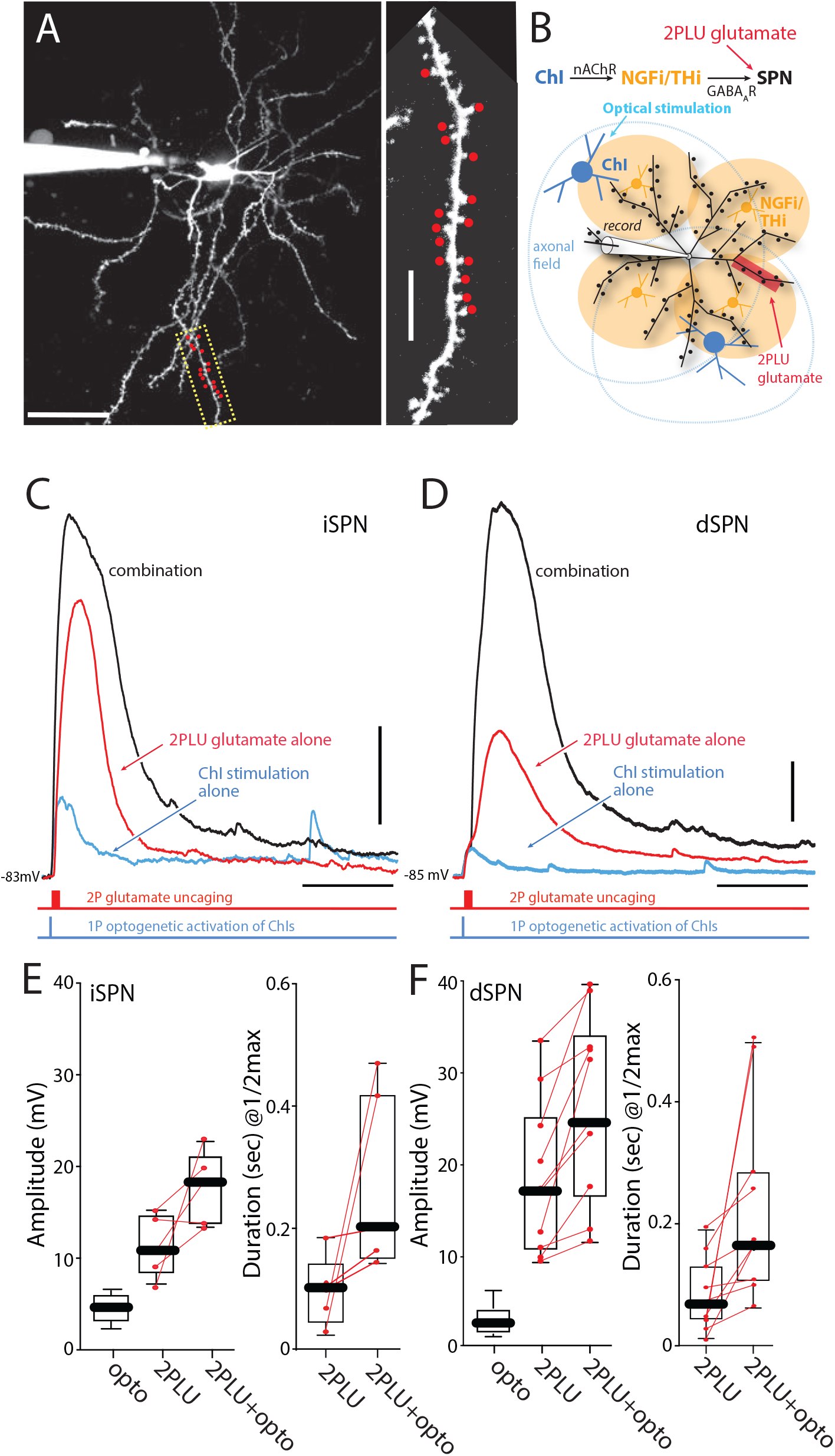
ChI evoked stimulation of NGF interneurons and synaptic GABA release enhances glutamate-evoked state transitions. **(A)** Maximum projection image of a visually identified dSPN from a D1R-tdTomato x ChAT-cre mouse with a high magnification image of a distal dendrite where 720nm 2-photon-laser-scanning microscopy spot uncaging of DNI-Glu (2PLU, 5 mM) was conducted (red dots). Tomato+ dSPNs were patched in whole-cell mode and the cells were loaded with Alexa 568 for clear identification of dendrites and spines. Scale bars = 40 µm cell, 5 µm dendrite. **(B)** Scheme for interrogating endogenous GABA release from NGFIs onto SPNs via optogenetic stimulation of ChAT-cre mice expressing Chronos. **(C)** Throughout the dendrites, glutamate uPSPs in dSPNs and iSPNs can be evoked by uncaging DNI-Glu (5 mM, 1 × 15 spines, 1 ms pulses at 500 Hz, red traces, 720 nm laser) while stimulating GABA release from NGFIs with the blue laser (1 × 3 ms pulse, blue traces, 473 nm, within ∼20 µm of the dendrite). **(D, E)** From the quiescent down-state, GABA_A_R activation is depolarizing and pushes SPNs toward enhanced dendritic integration in both dSPN and iSPN dendrites (Glu-2PLU + GABA_A_ opto = black trace, scale bars = 5 mV/200 ms). **(F, G)** Summary data showing the enhancement in amplitude and duration of the plateaus at ½ the maximum amplitude (1/2max) in iSPNs and dSPNs, respectively (iSPNs, Amp p=0.0040, 1/2max p=0.0312, n=5; dSPNs, Amp p=0.0020, 1/2max p=0.0034, n=10, Mann-Whitney U, one-tailed). All experiments are conducted in the appropriate cocktail of synaptic blockers: AP5 (50 µM), DNQX (5 µM), CGP-55845 (1 µM), MPEP (1 µM), and CPCCOEt (50 µM).

### Computational modeling of dendritic integration in SPNs

Although intriguing, the experimental results presented are limited by the inability to rapidly and precisely control the timing and location of GABAergic input to dendrites. Understanding how the timing and dendritic location of GABA_A_R activation modulates the response to clustered excitatory input could provide insight into the role of GABAergic interneurons in striatal computation. To help achieve a better grasp of the mechanisms underlying this interaction, a slightly modified NEURON model of a dSPN [13,27–29] was used to assess the impact of timing and location of GABAergic input on the response to clustered glutamatergic synaptic input to a stretch of distal dendrite. As observed experimentally, clustered glutamatergic input was able to generate NMDAR-dependent, dendritic spikes or plateau potentials when delivered to distal dendrites of a quiescent neuron (**Fig. 5A-C**).

**Figure 5.**
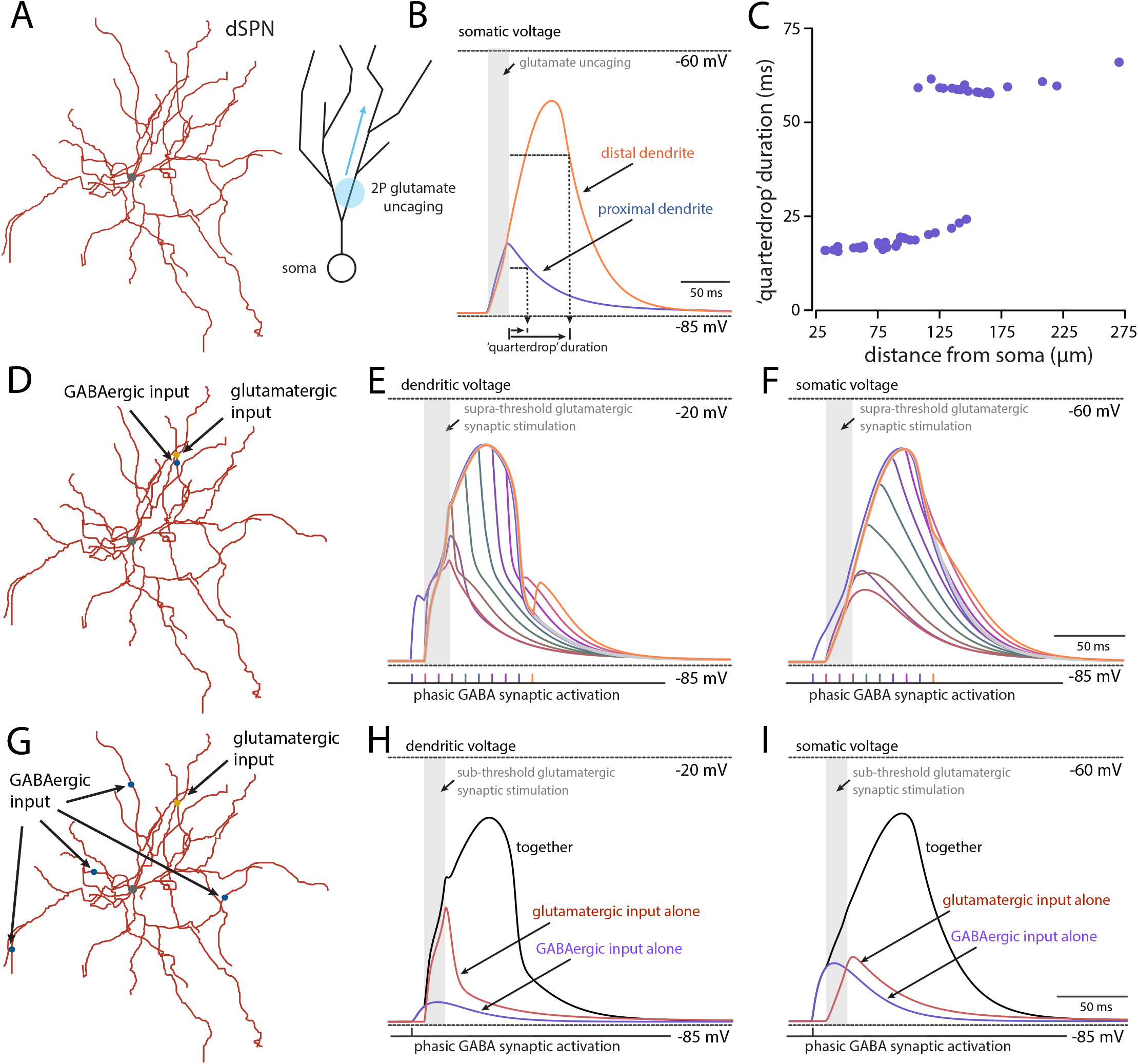
Computational modeling of dSPN dendritic activity. **(A)** Morphology of reconstructed dSPN with cartoon to illustrate the direction of stimulation along a dendrite. **(B)** Synaptic potentials recorded at the soma in response to clustered spine activation (18 neighboring glutamatergic synapses stimulated sequentially at 1ms intervals) in two separate dendrites. An up-state is generated in the distal (orange trace) but not proximal dendrite (blue trace). The quarterdrop duration is defined as the time interval between the last stimulation and the time for the membrane voltage to drop by one quarter of its peak value. **(C)** Quarterdrop interval (ms) plotted as a function of path distance from the center of that dendrite to the soma (µm) for every dendrite with spines (50) of the reconstructed dSPN. Up-state generation is only reliably observed in distal dendrites (>100 µm from cell soma). **(D)** Onsite phasic GABAergic activation and glutamatergic activation delivered to the same distal dendrite. Synaptic potentials recorded at the dendrite **(E)** and soma **(F)**. GABA synaptic activation comprised 3 simultaneous stimulations delivered 5 times at an interval of 1 ms to the midpoint of the dendrite. The timing of this phasic input was varied in a temporal manner relative to a fixed clustered supra-threshold glutamatergic input (delivered to 18 spines; 1 ms interval as before) in intervals of 10 ms from -10 (blue) to 80 ms (orange). For comparison, the effect of glutamatergic activation alone is illustrated by a thick gray line. Onsite GABAergic activation causes a dramatic cessation of dendritic and somatic potentials in a manner consistent with the relative timing of the two inputs. **(G)** Offsite phasic GABAergic activation delivered to 4 distal dendritic locations. A clustered glutamatergic input (15 spines; 1 ms interval) delivered to the same dendrite as before resulted in a subthreshold synaptic potential (red) at the dendritic site of delivery **(H)** and soma **(I)**. Similarly, the effect of only activating GABAergic synapses simultaneously at each of the four offsite dendritic locations (3 per dendrite; 12 in total) resulted in a moderate post-synaptic potential (blue trace). When delivered sequentially, with GABAergic activation preceding glutamatergic by 10 ms, the previously subthreshold glutamatergic input resulted in the generation of an up-state (black trace; *I* and *H*).

When GABAergic synapses were activated near glutamatergic synapses, the model behaved as previously described by Du et al. (2017). That is, GABAergic input at almost any point during the dendritic spike (when the local membrane potential was near -20 mV) led to inhibition of both the dendritic and somatic membrane potential (**Fig. 5D-F**). However, the impact of GABAergic input was very different when the site of stimulation was at some distance from that of glutamatergic stimulation. For example, if the GABAergic input was distributed at distal locations across the dendritic tree, as predicted to happen following ChI or NGFI/THI activation, the effect was consistently excitatory. To illustrate this point, the distributed GABAergic input was followed by a sub-threshold dendritic glutamatergic input. In this scenario, the combination of GABAergic and glutamatergic input led to a dendritic spike (**Fig. 5G-I**) – just as seen experimentally using optogenetic stimulation of GABAergic interneurons and 2P uncaging of glutamate.

In many of the previous studies examining the impact of GABA_A_Rs on dendritic integration of glutamatergic input, the focus has been on the role of timing and location dependent GABA_A_R - mediated shunting of iGluR-evoked EPSPs on the same dendrite [30–32]. As shown above, our results are largely consistent with this literature. Of particular interest is the timing dependence of the interaction. To explore this relationship in SPNs, NEURON simulations were run with glutamatergic and GABAergic input to distal dendrites (**Fig. 6A**). As expected, there was a strong timing dependence on the interaction between synaptic events. As experimentally shown by others [6,18], when the glutamatergic EPSP preceded a neighboring GABAergic input, the effect of opening GABA_A_Rs on the input impedance (i.e., shunting) was clearly evident (**Fig. 6B, C**).

**Figure 6.**
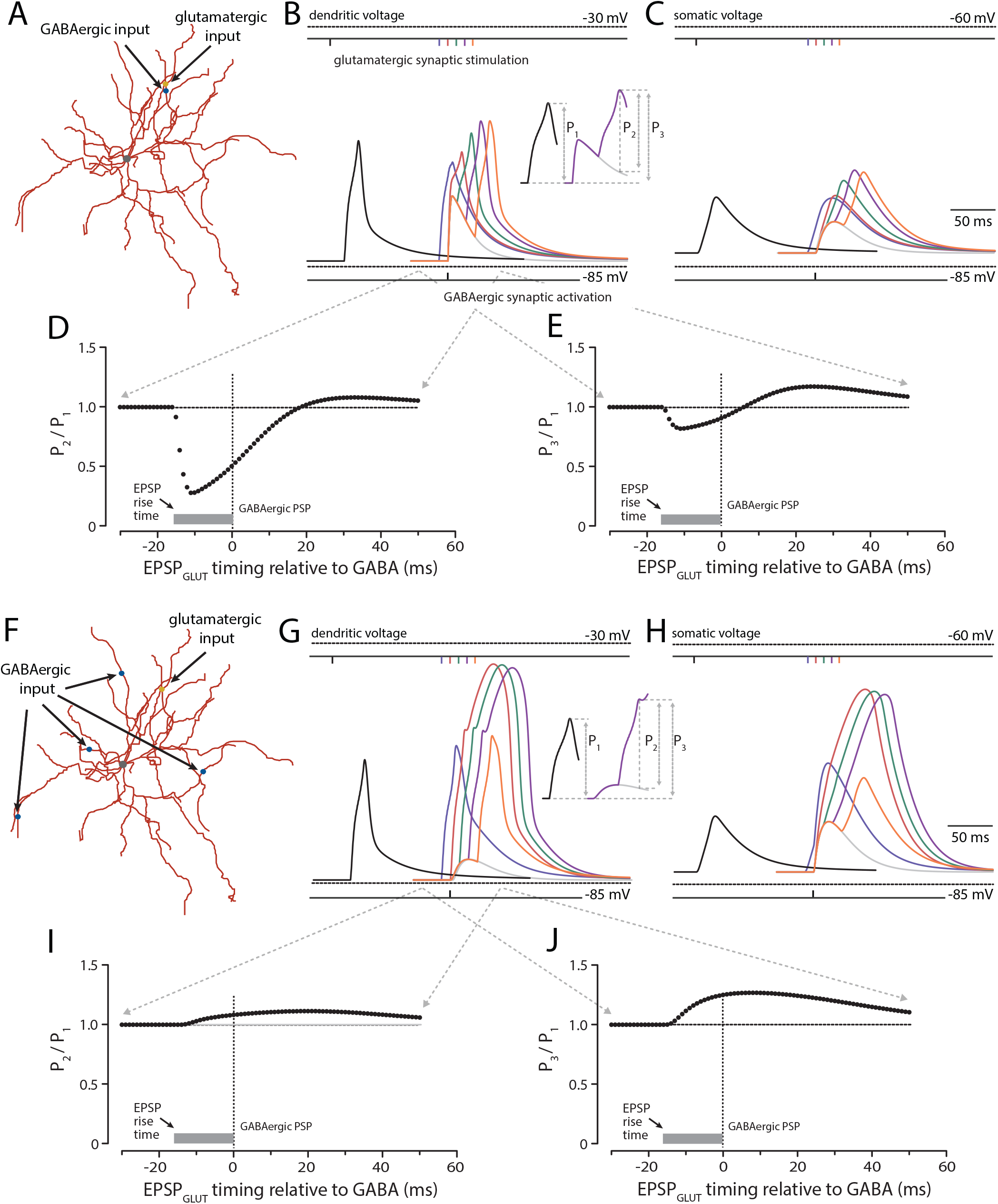
Temporal profile of interaction between glutamatergic and GABAergic synaptic activity. **(A)** On-site phasic GABAergic activity is delivered to the same dendrite as glutamatergic synaptic input. Synaptic potentials are illustrated at **(B)** dendritic site of glutamatergic activity and **(C)** cell soma. The black trace represents the effect of L-glutamate mediated excitation alone (15 neighboring spines activated at 1 ms intervals along the chosen dendrite). The light gray trace illustrates the effect of phasic GABAergic stimulation (3 simultaneous synaptic stimulations delivered 4 times at an interval of 1 ms to the same dendritic site giving a total of 12 GABA synapses activated). Color traces represent the effect of varying glutamatergic stimulation at temporal intervals relative to the fixed GABAergic input described (blue to orange illustrate 5 traces with ΔT = t_GLUT_ – t_GABA_ in the range of -10 to 30 ms, respectively at 10 ms intervals). Inset of *B* illustrates the measurement of P_1_, P_2_ and P_3_. The amplitude of the synaptic potential in the dendrite was measured in the absence of GABAergic activity (P_1_) or either relative to the amplitude of the depolarizing phasic GABAergic potential at the peak of the glutamatergic response (P_2_) or relative to the underlying baseline (P_3_). **(D)** and **(E)** show the sublinear effect of varying glutamatergic spine activation relative to a fixed onsite GABA synaptic input on P_2_ and P_3_ normalized to P_1_, respectively. **(F)** Off-site phasic GABAergic activity is delivered to 4 distal dendrites distinct from the dendrite receiving clustered spine excitation. As before, **(G)** and **(H)** show simulated synaptic potentials recorded at dendrite and soma. The black trace represents the same glutamatergic input as above (*i*.*e*., 15 spines activated at 1 ms intervals). The light gray trace represents the effect of phasic GABAergic activity delivered to the 4 distal dendrites at the same time (as 3 simultaneous synaptic simulations per dendrite giving a total of 12 GABA synapses activated). Again, as before, color traces show the effect of altering the timing of clustered spine activation relative to GABA activity. In contrast to on-site activity, ΔT = 0, 10 and 20 ms results in the generation of an up-state. Note that the peak of the EPSP_GLUT_ is still clearly visible (and measurable as P_2_) before the trailing up-state manifests. **(I)** and **(J)** illustrate the supralinear effect of varying glutamatergic spine activation relative to a fixed off-site GABA synaptic input on P_2_ and P_3_ normalized to P_1_, respectively.

However, when the glutamatergic input arrived later, there was synaptic summation, albeit sublinear at both dendritic (**Fig. 6B**) and somatic locations (**Fig. 6C**). To better illustrate the quantitative interaction between the two inputs at the dendritic site of stimulation, two plots were generated. In one, relative amplitude of glutamatergic EPSP (P2) with a concomitant GABAergic input was divided by the amplitude of the glutamatergic EPSP alone (P1) and then plotted as a function of the relative timing of the two inputs (**Fig. 6D**). This ratio (P2/P1) fell when the glutamatergic input preceded the GABAergic input and then rose when it trailed the GABAergic input. Similarly, if the ratio of the peak amplitude of the aggregate potential (P3) was divided by the peak amplitude of the isolated glutamatergic EPSP (P1) and plotted as a function of the relative timing of the two inputs, the ratio fell when the glutamatergic input preceded the GABAergic input, but then rose above 1 when the glutamatergic trailed the GABAergic input (**Fig. 6E**).

Of greater interest, given the architecture of striatal circuits, was how a diffuse GABAergic input, which should mimic conditions produced by ChI activation, would affect the interaction between synaptic events. To explore this interaction, the temporal relationship between a focal glutamatergic input to a distal dendrite and a GABAergic input to four neighboring dendrites was systematically varied (**Fig. 6F**). Not surprisingly, in this situation there was no shunting and the two inputs summed at both the dendritic (**Fig. 6G**) and somatic (**Fig. 6H**) locations – regardless of relative timing. In fact, GABAergic input enhanced the ability of glutamatergic synapses to trigger a dendritic spike **(Fig. 6G, H)**. To probe the dendritic interaction, the relative amplitude of the glutamatergic EPSP (measured at the inflection point of the membrane potential when there was an up-state) (P2) was divided by the amplitude of the isolated glutamatergic EPSP (P1) and then plotted as a function of the relative timing of the two inputs. At all intervals, the ratio was greater than or equal to 1 (**Fig. 6I**). A qualitatively similar plot was obtained by computing the ratio of the peak amplitude of the mixed PSP (P3) divided by the glutamatergic EPSP amplitude (P1) as a function of relative timing of the two inputs (**Fig. 6J**).

To better illustrate the role of location in dictating the shunting effect of a GABAergic synapse, the dendritic voltage and input impedance at the distal dendritic site was computed for 0 to 25 GABAergic synapses. When the GABAergic input was near the dendritic measurement site (**Fig. 7A**), the dendritic depolarization progressively increased with the number of GABAergic synapses, but the input impedance fell in parallel (**Fig. 7B, C**). In contrast, when the GABAergic synapses were placed on neighboring dendrites (**Fig. 7D**), the dendritic depolarization grew with the number of synapses, but there was little local change in input impedance (**Fig. 7E, F**).

**Figure 7.**
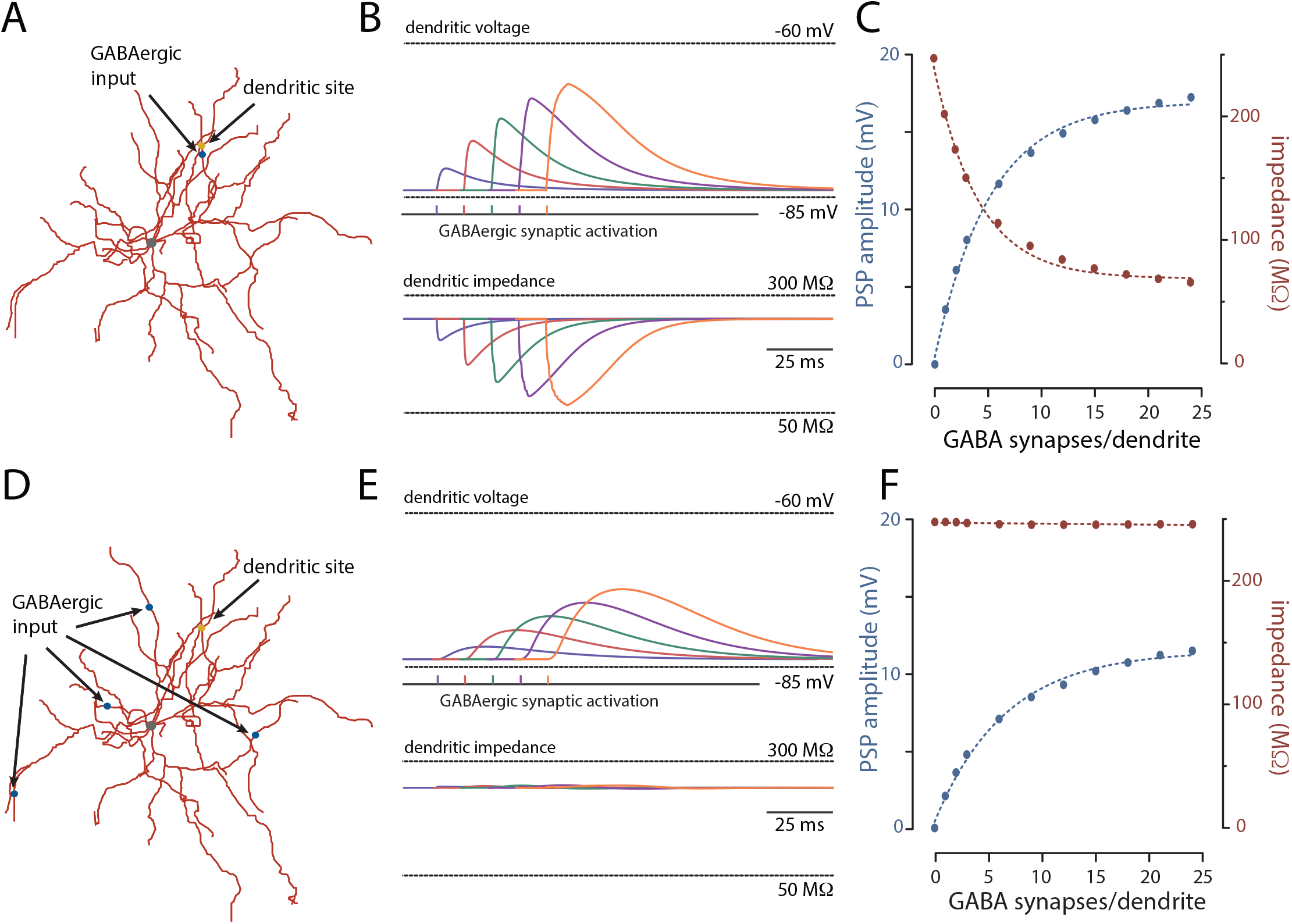
On vs off-site GABAergic activation and local input impedance. **(A)** On-site phasic GABAergic activity is delivered and recorded in the same dendrite. **(B)** Color traces (blue to orange; above) represent simulation GABAergic PSPs and local dendritic impedance (to 10 Hz; below) to increasing synaptic activation (3, 6… 15 GABA synapses). **(C)** Increasing PSP amplitude (blue) to GABAergic synaptic stimulation reduces local impedance (red). **(D)** Off-site phasic GABAergic activity is delivered to 4 distal dendrites and recorded in the same dendrite as in A. **(E)** and **(F)** as above. Note the lack of effect of increasing off-site GABAergic activity on local impedance.

The inference to be drawn from these simulations is that the interaction between dendritic glutamatergic and GABAergic synapses depends upon both location and timing. When co-localized, the timing of the two inputs dictates their interaction, as previously described [6,18]. However, when the GABAergic input is more diffuse, as predicted to occur with activation of ChIs or NGFIs, the relative timing of GABAergic and glutamatergic inputs becomes less important, and the two depolarizing inputs sum. This additivity allows GABAergic and glutamatergic synapses to work together to trigger dendritic spikes – as observed experimentally.

## Discussion

There are two main conclusions that can be drawn from the data presented. First, contrary to the developmental pattern described in many brain neurons, the GABA_A_R reversal potential in striatal SPNs does *not* undergo a significant hyperpolarizing shift with maturation [8,15]. The reversal potential of GABA_A_Rs in SPNs remained near -60 mV from weaning to adulthood.

Moreover, this relatively depolarized reversal potential was not attributable to NKCC1 expression, but rather to other factors like the expression of HCO_3-_/Cl^-^ co-transporters. Second, in resting SPNs engagement of striatal GABAergic interneurons produced a PSP much like that generated by iGluRs. Furthermore, ChI-mediated activation of striatal GABAergic interneurons that diffusely target dendrites enhanced the depolarization evoked by focused dendritic glutamatergic synaptic activity. In so doing, temporally aligned GABAergic input could sum and lower the threshold for the generation of dendritic spikes in response to glutamatergic signals. Taken together, these results argue that SPN GABA_A_R signaling should be considered as state-dependent and not strictly inhibitory or excitatory.

### The role of Cl-transporters in determining the GABA_A_R reversal potential in SPNs

A developmental shift in the reversal potential of GABA_A_Rs is seen in many neurons [8,15]. The initial experiments aimed at gaining a better understanding of this shift used a combination of pharmacological approaches and tissue RNA profiling to determine what was changing. These assays clearly pointed to a role for NKCC1 in pushing the GABA_A_R reversal potential to near spike threshold as tissue expression of NKCC1 was high in early stages of development and then declined, while KCC2 rose with age – paralleling the loss of ‘excitatory’ GABAergic signaling and the rise of ‘inhibitory’ signal. But these studies did not demonstrate cellular specificity of the shift in NKCC1/KCC2 expression. Subsequent work in the hippocampal glutamatergic neurons has partially filled this gap [33], but the generality of the model remains to be determined [34].

In contrast to the situation in many neurons studied to date, the SPN GABA_A_R reversal potential remained at a relatively depolarized value throughout development. Based upon the inferences drawn from previous work, our prediction was that this property of GABA_A_Rs in SPNs was attributable to the sustained expression of NKCC1 throughout development. However, in mRNA specifically harvested from iSPNs (using the RiboTag method), NKCC1 expression was essentially at the noise level in cells from both young and adult mice, whereas KCC2 expression was high throughout. The low level of NKCC1 expression in iSPNs was not due to our detection methodology, as at the tissue level, NKCC1 mRNA was readily seen. Although it is possible that SPNs expressed NKCC1 at embryonic or pre-weaning stages of development, NKCC1 is not a determinant of SPN GABA_A_R function in post-weaning mice.

What then is the explanation for the depolarized reversal potential of GABA_A_Rs in SPNs? A definitive answer to this question will require additional work, but our study does provide some clues. First, SPNs express three different isoforms of CA, which catalyzes the conversion of CO_2_ to H^+^ and HCO3^-^ [8]. Second, SPNs expressed mRNA for HCO3^-^/Cl^-^ co-transporters (e.g., AE3 and NCBE), which allows intracellular HCO3^-^ to be exchanged for extracellular Cl^-^, elevating the intracellular [Cl^-^]. Lastly, inhibiting CAs with acetazolamide led to a significant negative shift in the GABA_A_R reversal potential in perforated patch recordings. These results show that intracellular HCO_3-_ contributes GABA_A_R reversal potential, either by promoting cytoplasmic loading of Cl^-^ (via co-transporters) or by directly permeating GABA_A_Rs [8]. Distinguishing these mechanisms will require better pharmacological or genetic tools.

One of the distinctive physiological features of SPNs is that at rest, constitutively active, inwardly rectifying Kir2 K^+^ channels control the transmembrane potential, pulling SPNs close to the K^+^ equilibrium potential near -90 mV – the so-called ‘down-state’ of SPNs [3,35]. As these channels are distributed throughout the somatodendritic membrane they also are a major determinant of local input resistance and dendritic electrotonic structure [3]. With depolarization intracellular Mg^2+^ ions and polyamines are swept into the channel pore blocking it [36] Thus, in dendritic regions of SPNs in the down-state, GABA_A_R activation will depolarize the membrane and, in so doing, shut off Kir2 channels. As the depolarization produced by transient opening of synaptically activated GABA_A_Rs outlasts the change in input impedance, dendritic GABA_A_R activation should not only depolarize dendrites but increase their input impedance.

Even in the case of tonic GABA signaling GABA_A_R opening, and Kir2 K^+^ channel block will counteract one another producing less of a change in dendritic input impedance than would have been predicted otherwise.

Although in principle, GABA_A_R signaling in SPN dendrites should enhance the ability of glutamatergic synapses to promote spike generation, this has never been directly tested with synaptic stimulation. Previous work on this interaction in *ex vivo* brain slices used intrasomatic current injection to demonstrate the interaction [6]. To fill this gap, optogenetic tools were used in conjunction with two-photon uncaging of glutamate at visualized dendritic spines.

Optogenetic stimulation of ChIs was used to engage intrastriatal GABAergic interneurons that synapse on SPN dendrites [19]. Indeed optogenetic stimulation of ChIs evoked a robust gabazine-sensitive PSP in SPNs recorded in perforated patch mode as did direct optogenetic stimulation of NPY-positive GABAergic interneurons. In both cases, the GABAergic PSP was considerably delayed and slower than those evoked by SPN collaterals or fast-spiking interneurons [4,21]; most likely, this reflects the large axonal arbor of ChIs and NGFIs and the resulting diffuse release of GABA over the SPN dendritic tree.

To probe the interaction between this diffuse GABAergic input and focal activation of glutamatergic synapses, two-photon laser scanning uncaging of glutamate along a parfocal stretch of distal dendrite was used in conjunction with optogenetic stimulation of ChIs.

Spatiotemporally convergent glutamatergic input to distal dendrites of SPNs are capable of triggering local spikes [11–13,22], as already described in pyramidal neurons [37,38]. Importantly, when a ChI-evoked GABAergic PSP was followed a few milliseconds later by dendritic uncaging of glutamate, the SPN response to glutamate was enhanced and often reached the threshold for a local dendritic spike. Thus, from the down-state, GABAergic and glutamatergic synapses worked in concert to drive dendritic depolarization of SPNs.

### Functional implications for intrastriatal circuitry

The striatum is composed almost entirely of GABAergic neurons, the exception being ChIs. Although there has been a great deal of speculation about the function of the intrastriatal circuitry, its precise role in goal-directed behavior and habit execution remains obscure. In part, this lack of clarity may stem from thinking about GABAergic signaling as being exclusively inhibitory. Our results, in alignment with several previous reports, argue that this narrow view should be broadened. Consider for a moment the role of the intrastriatal GABAergic circuitry controlled by ChIs through fast nAChRs. ChIs have been implicated in several basal ganglia functions, including the response to salient events, set-shifting and movement sequencing [39]. On the face of it, the robust and diffuse coupling of ChIs to SPNs through ‘inhibitory’ GABAergic interneurons makes no sense in any of these contexts. However, the recognition that ChI-driven GABAergic input to quiescent iSPNs and dSPNs works in concert with glutamatergic signals to promote dendritic depolarization and pull SPNs into an ‘up-state’ creates a much more rational framework. Thus, ChIs serve to rapidly bring SPNs ‘online’ and ready to respond to cortical and thalamic signals directing movement. In this context, it is worth noting that *in vivo*, SPNs reside well above the K^+^ equilibrium potential [40–42]; this ‘resting’ state appears to be dynamically controlled by synaptic input, which very well could be created by the GABAergic input arising from spontaneously active GABAergic interneurons and those driven by tonically active ChIs [4].

It is also worth considering the potential role of fast spiking GABAergic interneurons (FSIs). FSIs preferentially target the perisomatic region of SPNs [43] and are widely considered to be part of a fast, feedforward inhibitory circuit linking the striatum with motor cortices [44]. While there is no doubt that FSI input to a spiking SPN is inhibitory, in a quiescent SPN, FSI input should act in precisely the same way as dendritic input and push the membrane potential toward -60 mV. Acting in this way, phasic FSI input to SPNs should facilitate – not inhibit – the response to trailing glutamatergic input from cortical pyramidal neurons. Thus, the timing of signals becomes a critical determinant of whether they should be considered ‘excitatory’ or ‘inhibitory’; that is, GABAergic input to SPNs should not be blanketly considered inhibitory.

## Materials and Methods

### Animals

The following transgenic male and female mice were used: Adora2a-eGFP (C57BL/6J), RRID:MMRC_010541-UCD; Drd1-tdTomato (FVB), RRID:MMRRC_030512-UNC; Chat-cre (C57BL/6J), RRID:MMRRC_017269-UCD; NPY-cre, (C57BL/6J), RRID:MMRRC_034810-UCD; and Adora2a-cre (C57BL/6J), RRID:MMRRC_034744-UCD. Chat-cre and NPY-cre mice were backcrossed to Adora2a-eGFP and DRD1-tdTomato reporter lines in house and used with the approval of the Northwestern University Animal Care and Use Committee and in accordance with the National Institutes of Health (NIH) *Guide for the Care and Use of laboratory Animals*. Mice were group-housed with food and water ad libitum on a 12-hour light/dark cycle with temperatures of 65° to 75°F and 40 to 60% humidity.

### Stereotaxic surgery

An isoflurane precision vaporizer (Smiths Medical PM) was used to anesthetize mice. Mice were then placed on a stereotaxic frame (David Kopf Instruments), with a Cunningham adaptor (Harvard Apparatus) to maintain anesthesia delivery during surgery. The skull was exposed, and a small hole was drilled at the desired injection site. The following stereotaxic coordinates were used: Striatum, AP = +0.74, ML = -1.85, DV = -3.50. The Allen Mouse Brain Atlas, online version 1, 2008 (https://atlas.brain-map.org/) was used as a reference for the coordinates and generating diagrams. For each mouse, the distance between bregma and lambda was calculated and used to adjust the coordinates. For AAV injections, ∼500 nl of viral vector was delivered using a glass micropipette (Drummond Scientific) pulled with a P-97 glass puller (Sutter Instruments). Surgeries for electrophysiology experiments utilizing Chronos were performed unilaterally while surgeries for RiboTag tissue collection were executed bilaterally.

Electrophysiology experiments using Chronos were performed after at least 21 postoperative days; tissue collection for RiboTag was performed 10 days after injection.

### RiboTag profiling

AAVs for expression of RiboTag under a cre-dependent promoter (AAV5-hsyn-DIO-Rpl22l1-3Flag-2A-eGFP-WPRE, titers 2.24 × 10^13^ viral genomes/ml) were stereotaxically injected into the striatum in Adora2a-cre mice at P18 or p170, as described above. Ten days after injection, mice were sacrificed and the striatal tissue expressing RiboTag was dissected out using fluorescence microscopy and then frozen at − 80°C. RiboTag immunoprecipitation was carried out as previously described [(M. Heiman, R. Kulicke, R. J. Fenster, P. Greengard, N. Heintz, Cell type-specific mRNA purification by translating ribosome affinity purification (TRAP). *Nat. Protoc*. **9**, 1282–1291 (2014)]. Briefly, tissue was homogenized in cold homogenization buffer [50 mM tris (pH 7.4), 100 mM KCl, 10 mM MgCl_2_, 1 mM dithiothreitol, cycloheximide (100 μg/ml), protease inhibitors, recombinant ribonuclease (RNase) inhibitors, and 1% NP-40]. Homogenates were centrifuged at 10,000*g* for 10 min, and the supernatant was collected and precleared with protein G magnetic beads (Thermo Fisher Scientific) for 1 hour at 4°C, under constant rotation. Immunoprecipitations were carried out with anti-Flag magnetic beads (Sigma-Aldrich) at 4°C overnight with constant rotation, followed by four washes in high-salt buffer [50 mM tris (pH= 7.4), 350 mM KCl, 10 mM MgCl_2_, 1% NP-40, 1 mM dithiothreitol, and cycloheximide (100 μg/ml)]. RNA was extracted using RNeasy Micro RNA extraction kit (QIAGEN) according to the manufacturer’s instructions.

### Quantitative real-time PCR

RNA was extracted from the dissected striatal tissue using RNeasy mini kit (QIAGEN). cDNA was synthetized by using the SuperScript IV VILO Master Mix (Applied Biosystems) and preamplified for 10 cycles using TaqMan PreAmp Master Mix and pool of TaqMan Gene Expression Assays (Applied Biosystems). The resulting product was diluted and then used for PCR with the corresponding TaqMan Gene Expression Assay and TaqMan Fast Advanced Master Mix. Data were normalized to *Hprt* by the comparative CT (2-DDCT) method. TaqMan probes were used for PCR amplification of *Hprt*, Mm03024075_m1, Slc12a2 (NKCC1), Mm01265955_m1, Slc12a5 (KCC2), Mm00803929_m1, Slc4a3 (AE3) Mm00436654_g1 and Slc4a10 (NCBE) Mm00473827_m1. Experimental Ct values were normalized to *hprt* values using the following formula: ΔCt = Ct (*gene of interest*) − Ct (*hprt*). The final expression levels were shown as ΔCt values.

### *Ex vivo* slice preparation

Coronal or parasagittal slices (275 μm thickness) were obtained from mice ranging in age from 4 weeks to 9 months. Mice were acutely anesthetized with a mixture of ketamine (50 mg/kg) and xylazine (4.5 mg/kg) and perfused transcardially with oxygenated ice-cold saline (4°C) containing in mM: 125 NaCl, 3 KCl, 2.5 MgCl_2_, 0.5 CaCl_2_, 25 NaHCO_3_, 1.25 NaH_2_PO_4_ and 10 glucose (saturated with 95% O_2_-5% CO_2_; pH 7.4; 300 mOsM/l). After perfusion, mice were decapitated, and the brains were rapidly removed. Slices were obtained in oxygenated ice-cold saline using a vibratome (VT1000S, Leica Microsystems). Slices were transferred to an ACSF-filled holding chamber containing in mM: 125 NaCl, 3 KCl, 1 MgCl_2_, 2 CaCl_2_, 25 NaHCO_3_, 1.25 NaH_2_PO_4_ and 10 glucose (saturated with 95% O_2_-5% CO_2_; pH 7.4; 300 mOsm/l) and held there for ∼30 min at 34° before being allowed to come to room temperature (21–25°C) where they remained until recording.

### Electrophysiological recordings

For electrophysiological recordings slices were transferred to a submersion-style recording chamber mounted on an Olympus BX51 upright microscope (60X/1.0 NA objective) equipped with infrared differential interference contrast. Whole-cell and perforated-patch clamp electrophysiological recordings were performed with Multiclamp 700B amplifier. Signals were filtered at 1KHz. Stimulation and display of electrophysiological recordings were obtained with custom-written freeware *WinFluor* (John Dempster, Strathclyde University, Glasgow, UK). http://spider.science.strath.ac.uk/sipbs/software_winfluor.htm) that synchronizes two-photon imaging and electrophysiology. Targeted electrophysiological recordings were obtained from visually identified iSPNs or dSPNs. Patch pipettes (4-4.5 MΩ) were prepared with a Sutter Instruments horizontal puller using borosilicate glass with filament and filled with (in mM): 120 potassium-D-gluconate, 13 KCl, 10 HEPES, 0.05 EGTA, 4 ATP-Mg_2_, 0.5 GTP-Na, 10 phosphocreatine-di (tris); pH was adjusted to 7.25 with KOH and osmolarity to 275-280 mOsm. In perforated-patch experiments, 10 μM gramicidin was added to the internal recording solution to induce chloride-impermeable pore formation along with 25 μM Alexa Fluor 568 hydrazide Na^+^ salt (Invitrogen) to visualize any potential pore rupture. All perforated-patch recordings were corrected for liquid junction potential. Electrophysiological characterization of neurons was made in current clamp configuration. The amplifier bridge circuit was adjusted to compensate for electrode resistance. Access resistances were continuously monitored, and experiments were discarded if changes >20% were observed. Digitized data were imported for analysis with commercial software (IGOR Pro 6.0, WaveMetrics, Oregon).

### Optogenetic blue light stimulation

Simultaneous electrophysiological and Chronos optogenetic photo-stimulation or RUBI-GABA uncaging were performed with a targeted focal spot blue laser (473 nm Aurora laser launch, Prairie Technologies) system using the *Photostimulus Editor* in *WinFluor*. The Point Photo-activation module (Prairie Technologies) allows two different stimulation areas, and intensities, with sub-μm (small spot) critical illumination or an additional lens to stimulate ∼8 μm (large spot) diameter photo-stimulation in the sample focal plane with the 60x/1.0 objective. To control the release of synaptic GABA, Chat-cre and NPY-cre mice were injected with Chronos (AAV-/hsyn-flex-chronos-GFP or AAV-/hsyn-flex-chronos-tdTomato; UNC GTC Vector Core) as described in the Stereotaxic Surgery section of these methods. To activate NPY or Chat Chronos containing axons in the striatum, the targeted 473 nm spots were positioned adjacent to individual dendritic spines to photo-stimulate presynaptic terminals impinging on iSPNs or dSPNs. The laser power was calibrated to evoke a somatic postsynaptic potential of 2-5 mV. Although the laser was aimed peri-dendritically, the blue excitation laser light will travel in a focusing hourglass (small spot) or column (larger spot) through the slice with likely activation of the large dendritic fields of the striatal interneurons, above and possibly below the sample focal plane, resulting in diffuse synaptic GABA_A_R activation of postsynaptic receptors. For simultaneous stimulation of 5-10 spines and the reversal potential experiments, the larger blue laser spot was used. Additionally, whole-field photo-stimulation through the 60X objective (26.5FN with ∼440 µm diameter exposure) was coordinated with an epi-fluorescence-based LED (475/30 nm, pE-100, CoolLED) reflected through an eGFP filter cube and controlled with the *Stimulus Editor* in *WinFluor*. The time synchronized results were displayed in the *WinFluor* main *Record Images and Signals* window.

### vTwo-photon Excitation Uncaging of DNI-Glutamate

Simultaneous two-photon laser uncaging (720nm) and optogenetic stimulation of synaptic GABA (473nm) were performed using a laser scanning microscope system (Ultima, Bruker Technologies; formerly Prairie Technologies) with a tunable imaging laser (Chameleon-Ultra1, Coherent Laser Group, Santa Clara, CA) and Olympus BX-51WI upright microscope with 60X/1.0NA water-dipping objective lens was used to locate and acquire a whole-cell patch clamp. 810 nm from the imaging 2P laser excited Alexa Fluor 568 (580-630 nm; R3896 PMT, Hamamatsu) to visualize dendrites of the patched soma and distal (>100 μm) dendritic spines from planar sections (∼20 µm) of the same dendrite with image zoom 4 and 50 µm FOV. Custom written software (*WinFluor*, John Dempster and its *PhotoStimulusEditor* module, Nicholas Schwarz; features now available in *PrairieView* 5.x) was used to direct, control, test, synchronize, and display electrophysiological recordings combined with laser imaging and photo-stimulation. Simultaneous 2PLU (720 nm, Coherent Chameleon) and single photon optogenetic stimulation of synaptic GABA (473 nm, Prairie Aurora Laser Launch) were provided by a second, separate, independently controlled galvanometer mirror pair in the Ultima system. The three laser beams were optically combined (760DCLPXR, Chroma Technologies) in the scan head and aligned to the microscope optical path. DNI-glutamate (5 mM, Femtonics, Budapest, Hungary) was perfused in the recorded area and then excited by the 720 nm 2P laser. Pulses of 1 ms duration (∼10 mW sample power) were delivered to single spines located in the same focal plane where the laser average power or spot location was calibrated to evoke a somatic excitatory postsynaptic potential of 1-2 mV for each spine. During synchronized acquisitions the blue laser GABA photo-stimulation (one pulse, 3ms duration) preceded the glutamate uncaging of ∼15 spines with 1 ms duration and 1 ms inter-stimulation interval. These experiments were all conducted in the appropriate cocktail of synaptic blockers: AP5 (50 µM), NBQX (5 µM), CGP-55845 (1 µM), MPEP (1 µM), and CPCCOEt (50 µM).

### Pharmacological reagents

Stock solutions were prepared before experiments and added to the perfusion solution or focally applied with pressure ejection in the final concentration indicated. Two-photon laser uncaging and optogenetic experiments were performed in the presence of AP5 (50 µM), NBQX (5 µM), CGP-55845 (1 µM), MPEP (1 µM), and CPCCOEt (50 µM). All drugs were obtained from Hello Bio, Sigma or Tocris.

### Confocal imaging

Fixed tissue was prepared by transcardially perfusing terminally anesthetized mice with phosphate-buffered saline (PBS; Sigma-Aldrich) immediately followed by 4% paraformaldehyde (PFA; diluted in PBS from a 16% stock solution; Electron Microscopy Sciences). The brain was then removed and transferred into PFA solution overnight before being thoroughly rinsed and stored in PBS at 4°C. Fixed brains were then sectioned into 50-μm-thick coronal slices on a Leica VT1200S vibratome and collected in PBS. The sections were positioned on microscopy slides (VWR), allowed to dry and mounted with ProLong Diamond (Thermo Fisher Scientific) and #1.5 glass coverslips (VWR). Mounted sections were stored at 4°C until imaged with an Olympus FV10i-DUC confocal laser scanning microscope, using 10×/0.4 (air) or 60×/1.35 (oil) objective.

FIJI (NIH) was used to adjust images for brightness, contrast and pseudo-coloring.

### Model

The NEURON + Python (Neuron 8.2; Hines & Carnevale, 1997) model of a morphologically reconstructed SPN was integrated into a previously established model [13,27-28]. Cytoplasmic resistivity (Ra) was set to 200 Ω cm and specific capacitance was 1µF cm^-2^. The compartmentalized model (https://senselab.med.yale.edu/ModelDB/ShowModel?model=266775&file=/lib/params_dMSN.json#tabs-2) is biophysically-detailed and comprised a total of 700 segments with the following active and passive conductance’s: transient fast inactivating Na^+^ (Naf), persistent Na^+^ (Nap), fast A-type K^+^ (Kaf), slowly inactivating K^+^ (Kas), inwardly-rectifying K^+^ (Kir), delayed rectifier K^+^ (Kdr), small conductance Ca^2+^-activated K^+^ (SK), large conductance Ca^2+^-activated K^+^ (BK), L-type Ca^2+^ (Ca_v_ 1.2 and 1.3), N-type Ca^2+^ (Ca_v_ 2.2), R-type Ca^2+^ (Ca_v_ 2.3) T-type Ca^2+^ (Ca_v_ 3.2 and 3.3).

Channel distributions over cellular compartments were as previously described and are presented in [supplementary table based on Table 2; Lindroos et al., 2018]. Synaptic spines were added to all dendritic locations further than 30 µm from the center of the cell soma. Spines were added at a density of 1.468 per µm to give a total of 5500 spines for the reconstructed dSPN. The spines comprised a cylindrical head with a diameter of 0.5 µm connected to dendrites via a neck 1 µm long with diameter 0.1 µm. The morphologically reconstructed model dSPN had a resting membrane potential of -84 mV; a modest hyperpolarizing current step of 200 pA gave a ‘rectified range’ input resistance of approximately 85 MΩ and membrane time constant of 10.5 ms. The dSPN had an estimated whole cell capacitance of 180 pF. Candidate spines were selected as separate nearest neighbors along a dendrite at a start point of approximately two-thirds the length. NMDA and AMPA conductances were inserted into spines to be activated. Synaptic currents were modelled using a two-state kinetic model where the normalized peak conductance is determined by rise and decay time constants T_1_ and T_2_ (T_2_ > T_1_) as per (Du et al., 2017; Lindroos et al., 2018, Lindroos & Kotaleski, 2020). The maximal conductances of AMPA and NMDA responses were 350 and 787.5 pS, respectively. The reversal potentials for AMPA and NMDA was 0 mV. Spines were activated at 1 ms intervals in succession with stimulation moving away from the soma as per electrophysiological activation. The threshold for generating an up-state in the absence of any GABAergic activation was 15 glutamatergic inputs. GABA synapses were inserted directly onto the same dendritic location (the midpoint of the chosen dendrite). The maximal conductance was 1000 pS and reversal potential set to -60 mV. GABAergic responses were generated by activating these synapses simultaneously in groups of 3 at an interval of 1 ms. So, for onsite activation, a total of 12 GABA synapses were activated. For offsite activation, 4 distal locations were selected with each activated at 3 GABA synapses simultaneously (*i*.*e*., simultaneously within and between the 4 chosen dendrites).

### Statistical Analysis

Data was graphically presented using non-parametric box and whisker plots. In these plots, the center line is the median, the edges of the box mark the interquartiles of the distribution and the lines extend to the limiting values of the sample distribution; outliers (defined as values further away from the median than 1.5 x Interquartile range) are marked as asterisks. Statistical significance was determined using non-parametric tests running on GraphPad Prism Version 9.0 (GraphPad Software).

## Acknowledgements

We wish to thank Sasha Ulrich, YU Chen and Abdelhak Belmadani for expert technical assistance, George Augustine for the Clomeleon plasmid and Jun Ding for help in getting the NEURON modeling work started. This study was supported by the CHDI Foundation, the JPB Foundation, ASAP, and NIH (NS 34696).

---

**Table S1:** Max conductance of monovalent cation channels in the model (S cm^-2^)

| Name | axon | soma | initial dendrite | final dendrite | spine neck | spine head | dendritic distribution | dendritic variables |
| --- | --- | --- | --- | --- | --- | --- | --- | --- |
| NaF | 9 | 6 | 6.9150e-1 | 3.1342e-10 | 0 | 0 | sigmoidal | a<sub>4</sub>=0; a<sub>5</sub>=1; a<sub>6</sub>=50; a<sub>7</sub>=10; g<sub>max</sub>=0.7 |
| NaP | 0 | 7e-4 | 0 | 0 | 0 | 0 | NA | NA |
| KaF | 0 | 1.5e-1 | 8.9343e-2 | 6.0235e-2 | 0 | 0 | sigmoidal | a<sub>4</sub>=0.5; a<sub>5</sub>=0.25; a<sub>6</sub>=120; a<sub>7</sub>=30; g<sub>max</sub>=0.12 |
| KaS | 7e-3 | 1.6e-2 | 3.5896e-2 | 3.0e-3 | 0 | 0 | exponential | a<sub>4</sub>=0.25; a<sub>5</sub>=5; a<sub>6</sub>=0; a<sub>7</sub>=-10; g<sub>max</sub>=1.2e-2 |
| Kdr | 0 | 9.4e-4 | 7.4375e-4 | 1.7554e-4 | 0 | 0 | sigmoidal | a<sub>4</sub>=0.25; a<sub>5</sub>=1; a<sub>6</sub>=50; a<sub>7</sub>=30; g<sub>max</sub>=7e-4 |
| Kir | 0 | 1.2e-4 | 2.4e-4 | 2.4e-4 | 0 | 1e-7 | linear | g<sub>max</sub>=2.4e-4 |
| BK | 0 | 1.3e-4 | 1e-4 | 1e-4 | 0 | 0 | linear | g<sub>max</sub>=1e-4 |
| SK | 0 | 2e-5 | 2e-5 | 2e-5 | 0 | 2e-5 | linear | g<sub>max</sub>=2e-5 |
| e<sub>pas</sub> | 1.25e-5 | 1.25e-5 | 1.25e-5 | 1.25e-5 | 1.25e-5 | 1.25e-5 | linear | g<sub>max</sub>=1.25e-5 |
| I<sub>m</sub> | 1e-3 | 0 | 0 | 0 | 0 | 0 | NA | NA |

**Table S2:**
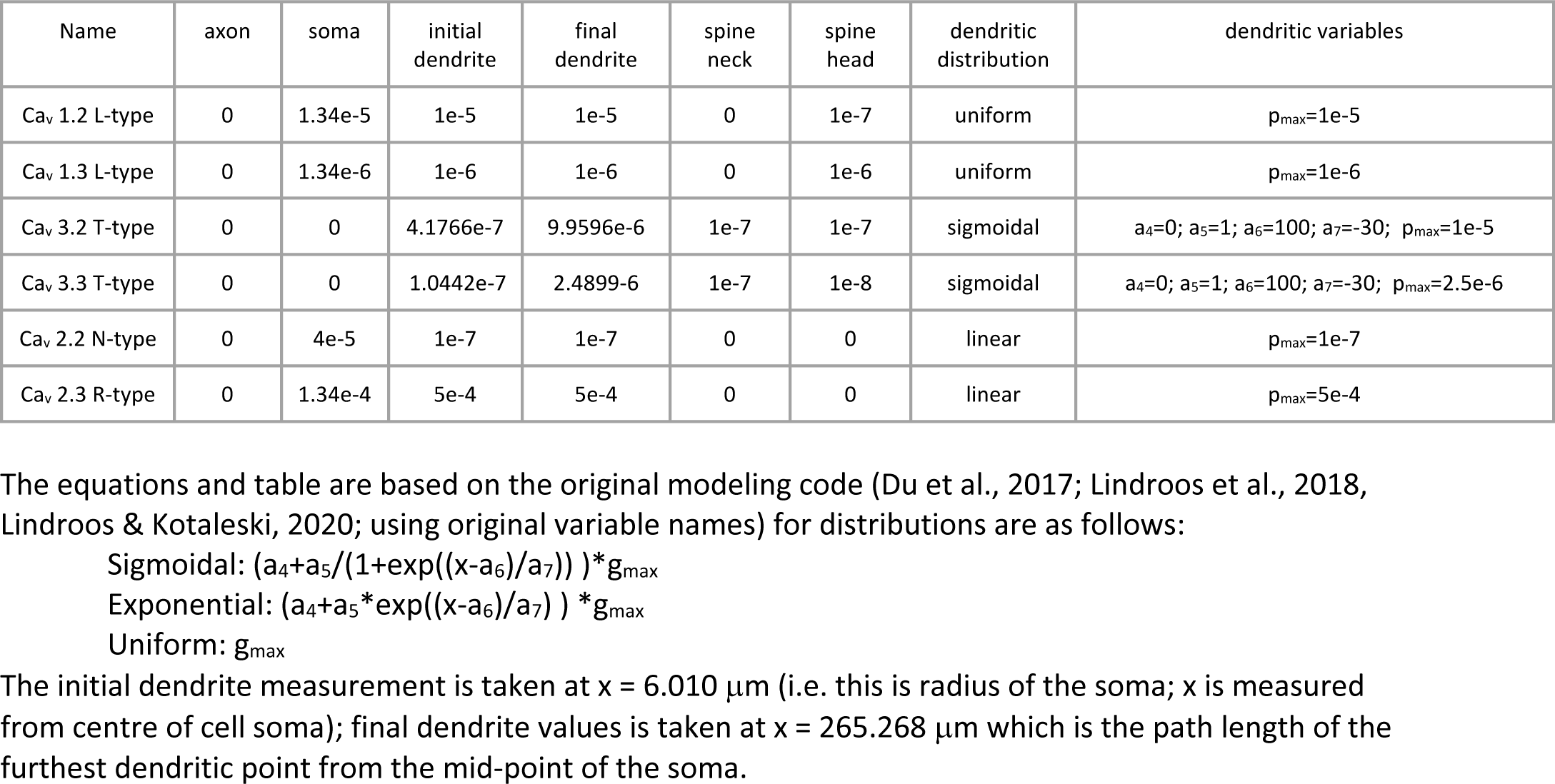
Max permeability of Ca2+ channels in model (cm s^-1^)

## Notes

### Competing Interest Statement

The authors have declared no competing interest.

### Summary of Updates

Figure 6 was corrected for typos

